# An economic dilemma between weapon systems may explain an arachno-atypical venom in wasp spiders (*Argiope bruennichi*)

**DOI:** 10.1101/2020.06.04.133660

**Authors:** Tim Lüddecke, Björn M. von Reumont, Frank Förster, André Billion, Thomas Timm, Günter Lochnit, Andreas Vilcinskas, Sarah Lemke

## Abstract

Spiders use venom to subdue their prey, but little is known about the diversity of venoms in different spider families. Given the limited data available for orb-weaver spiders (Araneidae) we selected the wasp spider *Argiope bruennichi* for detailed analysis. Our strategy combined a transcriptomics pipeline based on multiple assemblies with a dual proteomics workflow involving parallel mass spectrometry techniques and electrophoretic profiling. We found that the remarkably simple venom of *A. bruennichi* has an atypical composition compared to other spider venoms, prominently featuring members of the CAP superfamily and other, mostly high-molecular-weight proteins. We also detected a subset of potentially novel toxins similar to neuropeptides. We discuss the potential function of these proteins in the context of the unique hunting behavior of wasp spiders, which rely mostly on silk to trap their prey. We propose that the simplicity of the venom evolved to solve an economic dilemma between two competing yet metabolically expensive weapon systems. This study emphasizes the importance of cutting-edge methods to encompass smaller lineages of venomous species that have yet to be characterized in detail, allowing us to understand the biology of their venom systems and to mine this prolific resource for translational research.

## 1. Introduction

Spiders are diverse and species-rich arthropods that have conquered most terrestrial habitats. The 48,463 extant spider species (World Spider Catalog, 2020) share a common body plan that has changed little during ^~^380 million years of evolution. Spiders also possess a unique biochemical toolbox that uses a combination of venom and silk to subdue prey, contributing to their evolutionary success (Garrison et al., 2016). Moreover, they represent the only order of terrestrial animals in which almost all extant species feature a functional venom system and are thus considered as the most successful group of venomous animals (Saez et al., 2010). Accordingly, spiders play a pivotal ecological role as venomous predators by maintaining the equilibrium of insect populations (Riechert, 1974).

Venoms are complex mixtures of low-molecular-weight compounds, peptides and proteins, which act as toxins by disrupting important physiological processes when injected into prey (Fry et al., 2009; Nelsen et al., 2014). They are used for defense, predation or competitor deterrence, but in all cases they are physiologically expensive traits that have been optimized by strong selective pressure for specific functions. This evolutionary streamlining often results in high selectivity and target-specific bioactivity, meaning that animal venoms are now considered valuable bioresources in the field of drug discovery (Holford et al., 2018). Several blockbuster drugs have been derived from venom components (Holford et al., 2018), but they were also investigated as research tools, cosmetics, industrial enzymes or bioinsecticides (e.g. Duterte and Lewis, 2010; Pennington et al., 2018; Dongol et al., 2019; Saez and Herzig, 2019).

Spider venoms tend to be chemically more complex than other animal venoms and up to 1000 different venom components can be present in a single species (Herzig, 2019; Langenegger et al., 2019). It has been estimated, that the sum of all spider venoms could ultimately yield 10 million bioactive molecules, but only 0.02% of this diversity has been discovered thus far (Pineda et al., 2018; Lüddecke et al., 2019). Several promising drug candidates for stroke, pain, cancer and neural disorders have been identified in spider venoms (Osteen et al., 2016; Chassagnon et al., 2017; Richards et al., 2018; Wu et al., 2019). The major components of these spider venoms are low molecular weight inhibitor cysteine knot (ICK) peptides with robust tertiary structures conferred by the presence of a pseudoknot motif of interweaved disulfide bonds (Pallaghy et al., 1994; Norton and Pallaghy, 1998; Langenegger et al., 2019). Also described as knottins, such peptides are often neurotoxic and remarkably resistant to heat, osmotic stress and enzymatic digestion, making them ideal drug candidates (Langenegger et al., 2019). Although ICK peptides are found in other arthropod venoms (e.g. von Reumont et al., 2017; Drukewitz et al., 2018; Drukewitz et al., 2019; Özbek et al., 2019) the diversity of these peptides in spider venoms is unprecedented. Approximately 60% of all spider venom components accessible in UniProt (The UniProt Consortium, 2019) are ICK peptides, hence these peptides are commonly perceived as the principal component of spider venoms (e.g. Langenegger et al., 2019).

The analysis of venoms formerly relied on fractionation methods that require large amounts of starting material (e.g. Friedel and Nentwig, 1989; Hardy et al., 2013). Therefore, previous studies of spider venoms have focused on species with strong anthropocentric connections, such as those posing direct medical threats or those of extraordinary size, making the venom easier to access in sufficient quantities. This restricted analysis to members of the families Atracidae, Ctenidae, Theraphosidae, Sicariidae and Theridiidae, which represent a narrow sample of spider diversity (Lüddecke et al., 2019; World Spider Catalog, 2020). More recently, the advent of high-throughput methods compatible with miniscule samples has provided the means to expand the scope of such studies to less-accessible species (von Reumont, 2018; Herzig, 2019). Combinations of genomics, transcriptomics, proteomics and microbiomics (e.g. Oldrati et al., 2016; Ul-Hasan et al., 2019, Drukewitz & von Reumont 2019) now allow the analysis of venoms form previously neglected taxa (e.g. von Reumont et al., 2014) in an emerging field known as modern evolutionary venomics (von Reumont, 2018).

Several of the most species-rich spider lineages have not been studied at all in the context of venom systems (see Herzig et al., 2019 and Lüddecke et al., 2019). In this study, we therefore considered the family Araneidae (orb weavers), which is the third-largest spider family comprising 3078 extant species (World Spider Catalog, 2020). Orb weavers construct conspicuous and often large orb-like foraging webs, attracting the interest of evolutionary biologists (e.g. Bond et al., 2014; Fernandez et al., 2014). Little is known about their venoms and only one species (the Chinese orb weaver *Araneus ventricosus*) has been investigated using a venomics approach(Duan et al., 2013; Liu et al., 2016).

We selected the wasp spider *Argiope bruennichi*, which features a wasp-like banding pattern that may have evolved as a mimetic warning trait (Bush et al., 2008) This species is used as a model organism for the investigation of sexual dimorphism, chemical ecology, reproductive behavior, microbiome analysis and range expansion linked to climate change (e.g. Chinta et al., 2010; Welke and Schneider, 2010; Zimmer et al., 2012; Wilder and Schneider, 2017; Cory and Schneider, 2018; Ganske and Uhl, 2018; Sheffer et al., 2019; Krehenwinkel and Tautz, 2013; Krehenwinkel and Pekar, 2015; Krehenwinkel et al., 2015, Zhang et al., 2016). Its venom has been extracted for bioactivity assays but has not been analyzed in detail (Friedel and Nentwig, 1989). We applied a cutting-edge proteo-transcriptomics workflow, in which an automated multiple-assembly strategy is used as a first step to identify and annotate venom gland-specific transcripts. These are matched to proteome components detected directly using two parallel mass spectrometry (MS) platforms to achieve exhaustive and sensitive protein detection and identification. We discuss the functional components of the venom in the context of wasp spider hunting behavior, in which silk rather than a venomous bite is the primary weapon used to overpower prey (Eisner and Dean, 1976).

## 2. Materials and methods

### 2.1 Collection of specimens and sample preparation for transcriptomics and proteomics

Fourteen *A. bruennichi* adult females were collected in September 2018 in Gießen, Germany (N 50.5729555°/E 8.7280508°). Venom glands were dissected from CO_2_-anesthetized specimens under a stereomicroscope, washed in distilled water and submerged in phosphate-buffered saline (PBS). Venom was released by gentle compression with forceps and the extracts were centrifuged (10,000 × g, 10 min, room temperature) to pellet cell debris, before pooling the supernatants for lyophilization. The remaining venom gland tissue was transferred to in 1 ml RNAlater solution, pooled and stored at –80°C. Remaining body tissue was processed in the same manner.

### 2.2 Proteo-transcriptomics overall workflow

RNA-Seq was used to identify transcripts from venom glands and remaining body tissues, followed by assembly and automated annotation. Crude venom was analyzed by one-dimensional (1D) and twodimensional (2D) polyacrylamide electrophoresis (PAGE) before two parallel MS methods were used to identify the venom proteins. The first was matrix-assisted laser desorption ionization time of flight (MALDI-TOF)-MS to characterize intact peptides derived from the 2D gels, and the second was liquid chromatography electrospray ionization (nanoLC-ESI)-MS to characterize peptide fragments directly from the crude venom samples. Transcripts matching the proteins identified by MS were then analyzed in more detail by examining differential expression and annotations. The workflow is summarized in Figure 1.

**Figure 1:**
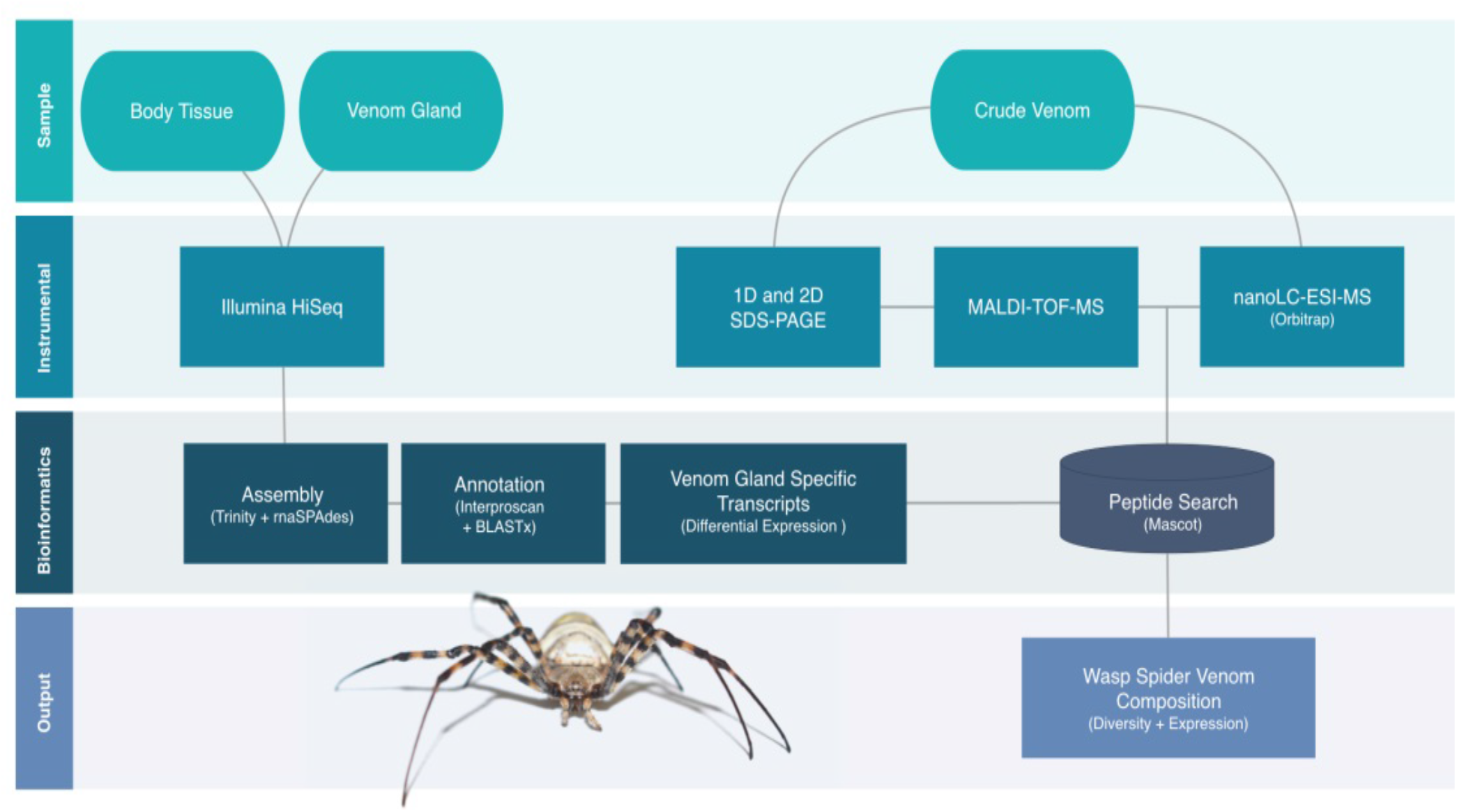
Proteo-transcriptomics workflow to characterize the venom of *A. brunnichi*. Transcriptomes of venom glands and body tissue were sequenced and assembled. Crude venom was analyzed by 1D/2D-PAGE before combinatorial MALDI-TOF-MS and nanoLC-ESI-MS. The final transcriptome assembly was used for the MS peptide search. Venom-specific transcripts matching detected proteins were then investigated in terms of expression levels and annotations.

### 2.3 Transcriptomics of venom gland and body tissue

#### 2.3.1 RNA extraction and sequencing

RNA extraction and sequencing was outsourced to Macrogen (Seoul, Korea). Following RNA extraction, libraries were constructed using the TruSeq RNA Sample Prep Kit v2 (paired-end, 151-bp read length). Quality was controlled by the verification of PCR-enriched fragment sizes on the Agilent Technologies 2100 Bioanalyzer with the DNA 1000 chip. The library quantity was determined by qPCR using the rapid library standard quantification solution and calculator (Roche). The libraries were sequenced on the Illumina HiSeq platform.

#### 2.3.2 Transcriptome assembly, annotation and quantification

Transcriptome data were processed using a modified version of our in-house assembly and annotation pipeline featuring different docker containers for enhanced reproducibility (Özbek et al. 2019). All containers (Table S1) were established using Nextflow v19.01.0 (https://www.nextflow.io/). Briefly, all input sequences were inspected using FastQC v0.11.7 before trimming in Trimmomatic v0.38 (Bolger et al., 2014; Andrews, 2019) using the settings 2:30:10, LEADING:5, TRAILING:5, SLIDINGWINDOW:4:15, and MINLEN:75. The trimmed reads were corrected using Rcorrector v1.0.3.1 (Song & Florea, 2015) and assembled *de novo* using a pipeline incorporating Trinity v2.8.4 (Grabherr et al., 2011; Haas et al., 2013) and rnaSPAdes v3.12 (Bushmanova et al., 2019) with and without error correction. All contigs were combined into a single assembly in which transcripts from all assemblers were merged if they were identical. The reads were remapped to the assembly using Hisat2 v2.1.0 (Kim et al., 2019) and expression values (transcripts per million, TPM) were calculated using stringtie v1.3.5 (Pertea et al., 2013; Pertea et al., 2016). SAM and BAM files were converted using Samtools v1.9 (Li et al., 2009). Open reading frames were then predicted with Transdecoder v5.0.2 (Haas et al., 2013) and annotated at the amino acid level using Interproscan v5.35-74 (Jones et al., 2014) and BLASTX v2.6.0+ (Camacho et al., 2009) searches against the Swissprot, Toxprot (The Uniprot Consortium, 2019) and Arachnoserver (Pineda et al., 2018) databases. The resulting assembly was used as a species-specific database for the identification of proteins detected by MS. Assemblies, sequencing raw data and transdecoder files are available at XXX Accession numbers to be added at manuscript acceptance XXX.

To avoid the overinterpretation of our data, differential expression analysis was applied to the two samples (venom gland versus remaining body tissue) and only putative venom components derived from transcripts with a logFC >2 within the venom gland dataset were considered further. Filtering steps were performed within the TBro v1.1.1 framework (Ankenbrand et al., 2016).

### 2.4 Venom proteomics

#### 2.4.1 Fractionation of venom proteins by PAGE

For 1D-PAGE, venom was mixed with tricine sample buffer (Bio-Rad) to make a total volume of 12 μl and incubated for 5 min at 95°C. The sample was then loaded onto a 16.5 % Mini-PROTEAN Tris-Tricine gel (Bio-Rad) in a Mini-PROTEAN Tetra System chamber (Bio-Rad) using 10x Tris-Tricine/SDS running buffer (Bio-Rad). Electrophoresis was carried out at 100 V for 100 min, and protein bands were detected with Flamingo stain (Bio-Rad).

For 2D-PAGE, contaminants were removed from the venom extract by the precipitation of 200 μg protein with 1:4 (v/v) chloroform/methanol (Wessel and Flugge, 1984). The protein pellet was redissolved in 260 μl lysis buffer (6 M urea, 2 M thiourea, 4% CHAPS, 30 mM DTT, and GE Healthcare 2% IPG buffer pH 3–10). GE Healthcare IEF strips (pH 3–10NL, 13 cm) were loaded with the sample by rehydration for 22 h and isoelectric focusing was carried out at gradients of 0–100 V/1 mA/2 W for 5 h, 100–3500 V/ 2 mA/5 W for 6 h, and 3500 V/2 mA/5 W for 6 h using a Multiphor II system (GE Healthcare). The IEF strip was then equilibrated for 15 min in 5 ml equilibration stock solution (6 M urea, 4% SDS, 0.01% bromophenol blue, 50 mM Tris-HCl pH 8.8, 30% (v/v) glycerol) containing 65 mM DTT, and then for 15 min in the same solution containing 200 mM iodacetamide. Proteins were separated in the second dimension on a 14% SDS polyacrylamide gel (Laemmli, 1970) in a Hoefer600 cell (GE Healthcare) for 15 min at 15 mA (100 V/15 W limits) and 4 h at 150 mA (400 V/60 W limits). The proteins were detected with Flamingo stain (Bio-Rad).

#### 2.4.2 MALDI-TOF-MS

The 2D gel was analyzed using PDQuest (Bio-Rad) and 152 spots (Table S2) were excised using the ExQuest Spot Cutter (Bio-Rad) and transferred into 96-well plates (Greiner Bio-One). The samples were digested simultaneously by using a MicroStarlet pipetting robot (Hamilton Robotics) to execute the following steps: the excised gel plugs were destained with 25 mM ammonium hydrogen carbonate containing 50% (v/v) acetonitrile, dehydrated with 100% acetonitrile, rehydrated in 50 mM ammonium hydrogen carbonate, dehydrated with 100% acetonitrile, dried at 56°C, rehydrated with 17 μl 25 mM ammonium hydrogen carbonate containing 4.5 ng/μl sequencing grade trypsin (Promega) and 0.025% Proteasemax (Promega) and incubated at 45°C for 2 h. Peptides were recovered by extraction with 15 μl 1% trifluoroacetic acid (Applied Biosystems) and stored at 4°C.

MALDI-TOF-MS was performed on an Ultraflex I TOF/TOF mass spectrometer (Bruker Daltonics) equipped with a nitrogen laser and a LIFT-MS/MS facility. Summed spectra consisting of 200–400 individual spectra were acquired in positive ion reflectron mode using 5 mg/ml 2,5-dihydroxybenzoic acid (Sigma-Aldrich) and 5 mg/ml methylendiphosphonic acid (Fluka) in 0.1% trifluoroacetic acid as the matrix. For data processing and instrument control, we used the Compass v1.4 software package consisting of FlexControl v3.4, FlexAnalysis v3.4, and BioTools v3.2. Data storage and database searches were carried out using ProteinScape v3.1 (Bruker Daltonics). Proteins were identified by Mascot v2.6.2 (Matrix Science) peptide mass fingerprinting using the venom gland transcriptome as a database. The search was restricted to peptides larger than 10 amino acids with a mass tolerance of 75 ppm. Carbamidomethylation of cysteine was considered as a global modification, the oxidation of methionine was considered as a variable modification, and one missed cleavage site was allowed. Only peptides with a Mascot Score > 80 were considered for further analysis (Table S3).

#### 2.4.3 NanoLC-ESI-MS

We dissolved 10 μg of protein were in 25 μl ammonium bicarbonate containing 0.1 ProteasMax. Cysteine residues were reduced with 5 mM DTT for 30 min at 50°C and modified with 10 mM iodacetamide for 30 min at 24°C. The reaction was quenched with excess cysteine before adding 0.025 ng/μl trypsin in a total volume of 100 μl. After incubation at 37°C for 16 h, the reaction was stopped by addition trifluoroacetic acid to a final concentration of 1 %. The sample was then purified using a C18-ZipTip (Millipore), dried under vacuum and redissolved in 10 μl 0.1 % trifluoroacetic acid.

For analysis, 1 μg of the sample was loaded onto a 50-cm μPAC C18 column (Pharma Fluidics) in 0.1% formic acid at 35°C. Peptides were eluted with a 3–44 % linear gradient of acetonitrile over 240 min followed by washing with 72% acetonitrile at a constant flow rate of 300 nl/min using a Thermo Fisher Scientific UltiMate 3000RSLCnano device. Eluted samples were injected into an Orbitrap Eclipse Tribrid MS (Thermo Fisher Scientific) in positive ionization mode via an Advion TriVersa NanoMate (Advion BioSciences) with a spray voltage of 1.5 kV and a source temperature at 250°C. Using data-independent acquisition mode, full MS scans were acquired every 3 s over a mass range of *m/z* 375–1500 with a resolution of 120,000 and auto-gain control (AGC) set to standard with a maximum injection time of 50 ms. In each cycle, the most intense ions (charge state 2–7) above a threshold ion count of 50,000 were selected with an isolation window of 1.6 *m/z* for higher-energy collisional (HCD) dissociation at a normalized collision energy of 30%. Fragment ion spectra were acquired in the linear ion trap with the scan rate set to rapid, a normal mass range, and a maximum injection time of 100 ms. Following fragmentation, selected precursor ions were excluded for 15 s.

Data were acquired with Xcalibur v4.3.73.11. (Thermo Fisher Scientific) and analyzed using Proteome Discoverer v2.4.0.305 (Thermo Fisher Scientific). Mascot v2.6.2 was used to search against the transcriptome database. A precursor ion mass tolerance of 10 ppm was applied. Carbamidomethylation of cysteine was considered as a global modification, the oxidation of methionine was considered as a variable modification, and one missed cleavage site was allowed. Fragment ion mass tolerance was set to 0.8 Da for the linear ion trap MS^2^ detection. The false discovery rate (FDR) for peptide identification was limited to 0.01 using a decoy database. For subsequent analysis, we only considered proteins with a Mascot Score > 30 and at least two verified peptides (Table S4). The raw proteomic raw data are available at PRIDE XXX Accession numbers to be added at manuscript acceptance XXX.

## 3. Results

### 3.1 The *A. bruennichi* venom gland and body tissue yield high-quality transcriptome libraries

Venom glands were dissected from 14 female *A. bruennichi* specimens and the venom was extracted and set aside for proteomic analysis. The venom glands and remaining body tissues were separately pooled, and RNA was extracted for RNA-Seq analysis. The resulting paired-end libraries were checked for DNA quantity and quality. The concentration of the venom gland transcriptome library was 116.26 ng/μl (fragment size = 387 bp) and the concentration of the remaining body tissue library was 91.47 ng/μl (fragment size = 363 bp). The venom gland transcriptome contained a total of 133,263,138 paired-end reads with a GC content of 42.2%, a Q20 of 98.2% and a Q30 of 94.5%. The remaining body tissue transcriptome contained a total of 145,808,360 paired-end reads with a GC content of 41.6%, a Q20 of 98.3% and a Q30 of 94.8%. The libraries were sequenced and annotated using our automated pipeline.

### 3.1 Only large *A. bruennichi* venom proteins are detected by SDS-PAGE and MALDI-TOF-MS

The crude venom set aside prior to RNA extraction was first fractionated by 1D-PAGE. The lanes representing both concentrations showed identical banding patterns after staining and the vast majority of the protein bands were in the 25–100 kDa range, with a few weaker bands of 15–25 kDa and no prominent bands below 15 kDa (Figure 2). To characterize the properties of these proteins in more detail, the venom sample was fractionated by 2D-PAGE. The isoelectric focusing step (pH 3–10) revealed proteins with a range of pI values, although predominantly focused around pH 7 (Figure 2). In agreement with the 1D-PAGE results, the orthogonal SDS-PAGE step indicated that most spots represented proteins of 25 kDa or more, with only a few in the size range 10–25 kDa and hardly any below 10 kDa (Figure 2).

**Figure 2:**
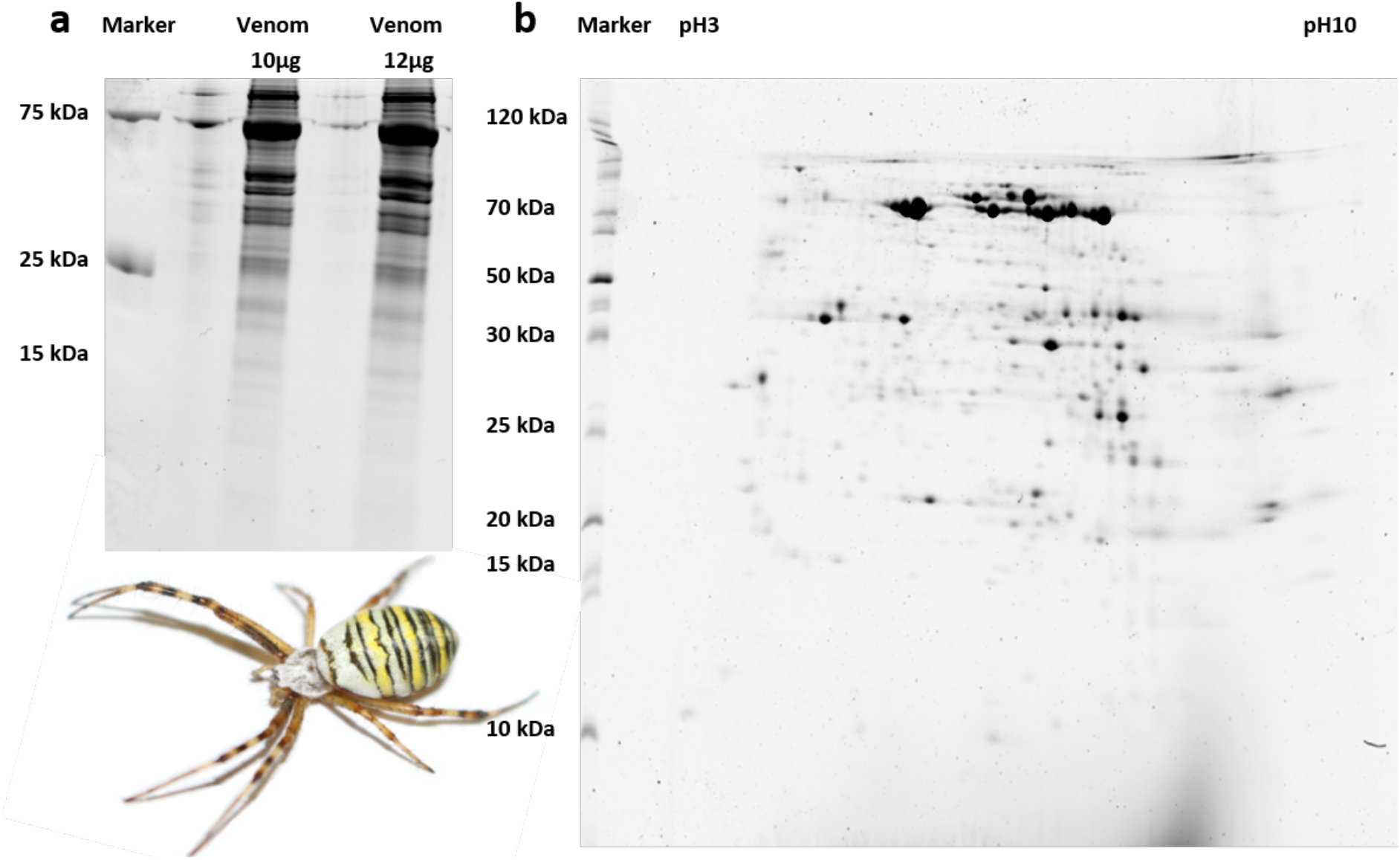
Analysis of *A. bruennichi* venom proteins by PAGE. (a) 1D-PAGE of venom proteins at two concentrations, showing identical banding patterns with most proteins larger than 25 kDa. (b) 2D-PAGE, showing that the proteins cover a range of pI values but cluster around pH 7, and conforming that most proteins are larger than 25 kDa.

We excised 152 spots from the 2D gels for MALDI-TOF-MS analysis, 41 of which matched to significantly enriched predicted coding regions from our venom gland transcriptome while the remainder matched non-venom proteins. Among the 41 venom-related spots, only six were ultimately assigned to protein classes known to be present in other animal venoms (Table 1). The sequences of the venom proteins identified in *A. bruennichi* were highly similar to the toxins previously identified its close relative, the Chinese orb-weaver *A. ventricosus* (Duan et al., 2013). Their molecular masses fell with the range 28.3–50.5 kDa and functional annotation revealed that they all belong to the cysteine-rich secretory protein, antigen 5 and pathogenesis-related protein 1 (CAP) superfamily.

**Table 1:** Identification of *A. bruennichi* venom proteins by MALDI-TOF-MS. Among 152 spots excised from 2D gels, 41 represented proteins that were enriched in the venom glands and six of these were similar to previously-identified venom components, all putative members of the CAP superfamily.

| Spot ID | Class | Score | kDa | Peptides | Coverage [ %] | ppm |
| --- | --- | --- | --- | --- | --- | --- |
| 8501 | CAP | 136.0 | 48.9 | 14 | 21.1 | 13.81 |
| 7309 | CAP | 70.7 | 28.3 | 6 | 18.3 | 7.42 |
| 7501 | CAP | 182.0 | 50.0 | 18 | 35.3 | 23.55 |
| 6502 | CAP | 85.9 | 50.5 | 10 | 24.8 | 22.95 |
| 6502 | CAP | 124.0 | 45.7 | 15 | 33.7 | 23.73 |
| 6502 | CAP | 89.2 | 48.1 | 11 | 25.3 | 23.57 |

CAP = cysteine-rich secretory protein, antigen 5 and pathogenesis-related protein 1

### 3.2 Further *A. bruennichi* venom proteins are revealed by high resolution nanoLC-ESI-MS

In a parallel proteomics workflow, the crude venom was analyzed by high-resolution nanoLC-ESI-MS (Orbitrap) revealing a total of 1806 protein groups, including 415 predicted coding regions matching significantly enriched predicted coding regions from our venom gland transcriptome. From this subset, we retrieved 54 protein groups with putative venom functions, representing 20 different protein families. Many of these protein families have previously been identified in spider venoms, including Kunitz-type serine protease inhibitors, prokineticin, EF-hand proteins, MIT-atracotoxins, astacin-like metalloproteases and ICK peptides. Three others showed similarities to hormones and neuropeptides (insulin-like growth factor binding protein (IGFBP), diuretic hormone and ITG-like peptides). Another class of proteins showed high sequence similarity to uncharacterized toxins previously isolated from *A. ventricosus* (Duan et al., 2013). BLAST searches did not recover any further similar sequences, so the remaining proteins were defined as “unidentified aranetoxins”. The nanoLC-ESI-MS experiment also confirmed our MALDI-TOF-MS data by showing that *A. bruennichi* venom contains multiple CAP proteins which are also the most abundant proteins among the venom components (Table 2).

**Table 2:** Identification of *A. bruennichi* venom proteins by nanoLC-ESI-MS. The analysis of peptide fragments allowed us to identify protein groups with putative venom functions, representing 20 different protein families. Confirming the parallel MALDI-TOF-MS analysis, most proteins could be assigned to the CAP superfamily.

| Protein Class | Matched Peptides | MW [kDa] | calc. pI | Mascot Score | Coverage |
| --- | --- | --- | --- | --- | --- |
|  |  |  |  |  | [ %] |
| Kunitz | 2 | 25.1 | 7.46 | 38 | 8 |
| Cystatin | 2 | 15.6 | 7.33 | 48 | 22 |
| Diuretic hormone-like | 2 | 14.6 | 9.55 | 267 | 23 |
| Techylectin | 3 | 40.3 | 7.17 | 88 | 7 |
| IGFBP | 2 | 18.8 | 4.92 | 63 | 16 |
| Venom protein 11 | 2 | 9.5 | 8.00 | 234 | 35 |
| 5' Nucleotidase | 4 | 24.3 | 4.54 | 228 | 17 |
| Prokineticin | 4 | 13.8 | 7.97 | 636 | 33 |
| Unclassified aranetoxins | 2 | 8.4 | 7.71 | 56 | 39 |
| Unclassified aranetoxins | 5 | 12.9 | 9.57 | 220 | 32 |
| Unclassified aranetoxins | 4 | 8.3 | 8.18 | 202 | 39 |
| Unclassified aranetoxins | 3 | 8.3 | 8.07 | 630 | 55 |
| Unclassified aranetoxins | 4 | 8.2 | 8.18 | 206 | 39 |
| MIT-atracotoxin | 4 | 10.7 | 4,94 | 448 | 56 |
| MIT-atracotoxin | 5 | 9.8 | 5.50 | 140 | 66 |
| Thyroglobulin-like | 3 | 10.9 | 7.56 | 134 | 31 |
| CAP | 16 | 48.9 | 8.44 | 1904 | 46 |
| CAP | 5 | 17.0 | 5.14 | 604 | 32 |
| CAP | 11 | 28.9 | 9.09 | 944 | 44 |
| CAP | 15 | 51.0 | 8.50 | 1127 | 40 |
| CAP | 9 | 28.3 | 7.66 | 1918 | 59 |
| CAP | 14 | 50.0 | 7.65 | 2164 | 40 |
| CAP | 8 | 50.5 | 7.97 | 1661 | 26 |
| CAP | 14 | 48.0 | 8.07 | 1020 | 46 |
| CAP | 3 | 25.1 | 6.84 | 173 | 27 |
| CAP | 23 | 51.2 | 7.77 | 3697 | 57 |
| CAP | 18 | 50.8 | 8.19 | 2333 | 52 |
| CAP | 6 | 13.2 | 8.79 | 311 | 49 |
| S10 peptidase | 4 | 51.4 | 8.07 | 118 | 15 |
| Leucine-rich-repeat domain | 20 | 39.7 | 4.93 | 4279 | 71 |
| Leucine-rich-repeat domain | 10 | 41.0 | 5.29 | 771 | 34 |
| Leucine-rich-repeat domain | 7 | 39.2 | 5.11 | 215 | 27 |
| Leucine-rich-repeat domain | 9 | 36.9 | 5.74 | 428 | 37 |
| Leucine-rich-repeat domain | 5 | 41.2 | 5.50 | 117 | 23 |
| EF-hand | 7 | 22.3 | 5.25 | 112 | 33 |
| ITG-like peptide | 10 | 26.6 | 4.78 | 1666 | 51 |
| ITG-like peptide | 8 | 24.7 | 4.96 | 673 | 38 |
| ICK | 8 | 15.0 | 6.15 | 1076 | 40 |
| ICK | 2 | 14.7 | 4.68 | 209 | 14 |
| ICK | 4 | 15.7 | 6.76 | 59 | 24 |
| Astacin-like metalloprotease | 2 | 14.4 | 5.87 | 76 | 13 |
| Astacin-like metalloprotease | 2 | 10.9 | 4.79 | 149 | 15 |
| Astacin-like metalloprotease | 3 | 17.0 | 7.93 | 50 | 24 |
| Putative chitinase | 2 | 18.0 | 5.19 | 112 | 11 |
| Putative chitinase | 3 | 47.4 | 11.15 | 53 | 13 |
| Putative chitinase | 8 | 30.1 | 6.49 | 219 | 39 |
| Putative chitinase | 12 | 35.5 | 7.30 | 1159 | 45 |
| Serine protease | 3 | 55.2 | 6.13 | 54 | 7 |
| Serine protease | 3 | 99.1 | 6.27 | 100 | 4 |
| Serine protease | 5 | 86.2 | 6.54 | 135 | 9 |
| Serine protease | 8 | 53.2 | 6.40 | 608 | 24 |
| Serine protease | 8 | 53.2 | 6.64 | 469 | 23 |

### 3.3 Data integration reveals that *A. brunnichii* venom is atypical for spiders

The transcriptomic and proteomic data were integrated for the comprehensive analysis of venom composition in terms of the diversity and abundance of venom proteins. In terms of overall diversity, the CAP superfamily was the most represented, with 15 different CAP proteins accounting for more than 27% of all the identified protein components. Leucine-rich repeat proteins and unclassified aranetoxins were also well represented, each with five members accounting for ^~^9% of the total diversity. We identified four putative chitinases and serine proteases, each accounting for ^~^7.5% of the total diversity, and three ICKs and astacin-like metalloproteases, each accounting for ^~^5.5% of the total diversity. There were two members of the MIT-atracotoxin and ITG-like peptide families, each contributing ^~^3.5% of the total diversity. Finally, the Kunitz serine protease inhibitor, cystatin, diuretic hormone-like peptide, techylectin, IGFBP, venom protein 11, 5’ nucleotidase, prokineticin, thyroglobulin, S10 peptidase and EF-hand families were represented by one member, each contributing <2% to the total diversity (Figure 3). In terms of abundance, CAPs represented 64.3% of the total protein content of *A*. *bruennichi* venom and were by far the most dominant component, followed by ITG-like peptides (9.5%), unclassified aranetoxins (7.7%) and leucine-rich repeat proteins (7.7%). The other components were expressed at much lower levels, with ICKs contributing only 3.3% of the total protein content, followed by putative chitinases (2.6%) and serine proteases (2.7%), and the others each contributing <1% (Figure 3).

**Figure 3:**
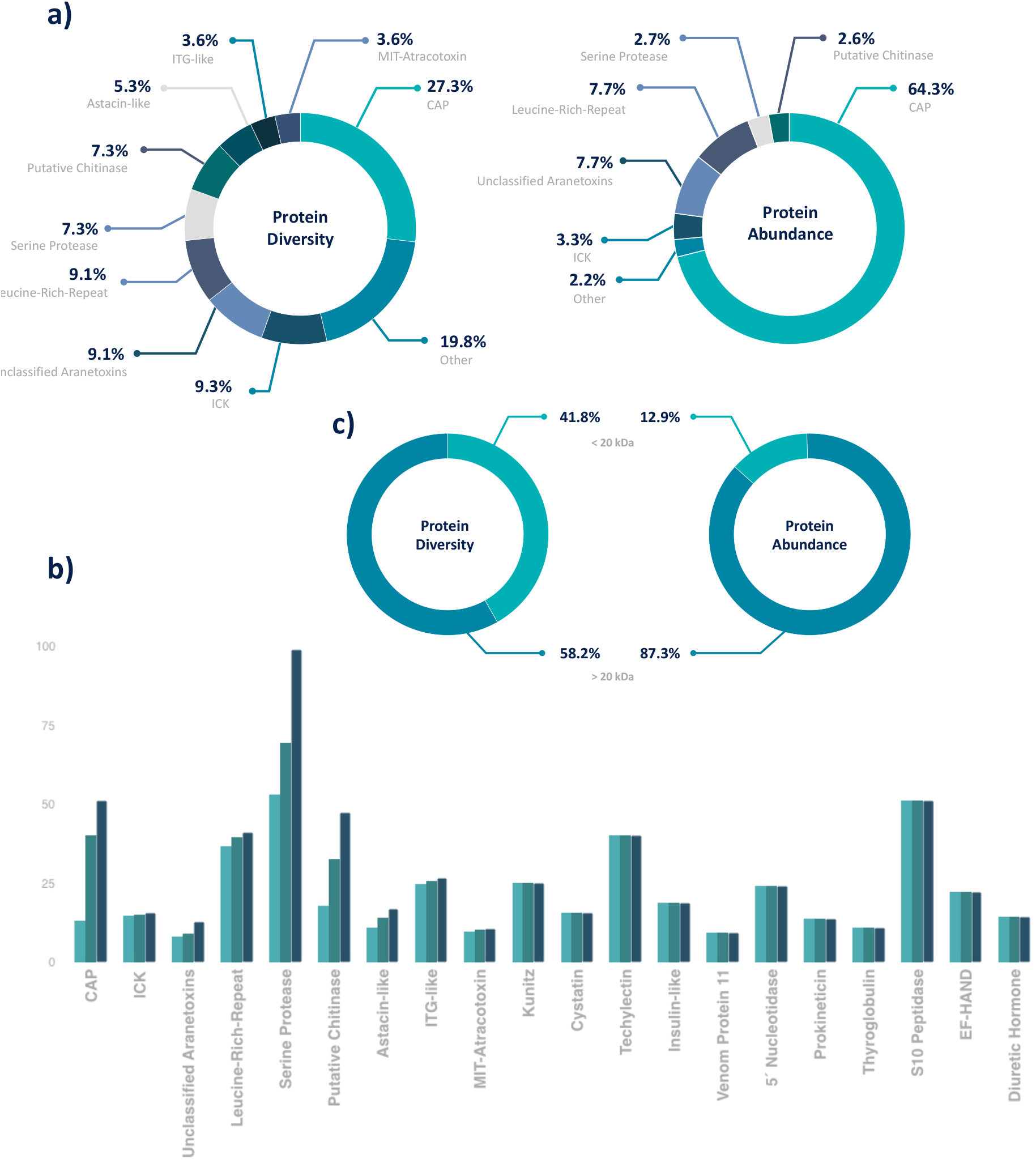
The venom protein profile of *A. bruennichi*. (a) Pie charts depict the venom composition in terms of *protein diversity* based on the number of distinct predicted coding sequences compared to *protein abundance* based on the transcripts per million reads for each coding sequence. By both measures, CAP family proteins are the dominant venom component, with 15 different members many expressed at high levels. (b) The molecular weight (kDa) of identified venom proteins, with the lowest, average and highest molecular weight per group from left to right. (c) Distribution of small (< 20 kDa) and large (> 20 kDa) proteins in terms of protein diversity and protein abundance.

In agreement with the 1D/2D-PAGE experiments, higher-molecular-weight proteins accounted for most of the diversity of the *A*. *bruennichi* venom proteome and are also the most abundant components. We identified 23 proteins/peptides with molecular masses < 20 kDa but these accounted for only ^~^42% of the proteome diversity and only ^~^13% of the total protein content (Figure 3).

## 4. Discussion

Spider venoms typically consist of mostly low-molecular-weight peptides, with ICK peptides as the predominant neurotoxic components. For example, ICK peptides represent 93% of the diversity in *Phoneutria nigriventer* venom (Diniz et al., 2018) and 42 of 46 identified venom components in the barychelid *Trittame loki* (Undheim et al., 2013). In *Cupiennius salei*, short cationic peptides and ICK peptides together comprise 39% of the venom components, whereas larger proteins only contribute 15% to its diversity (Kuhn-Nentwig et al., 2019). The predominance of ICK peptides has also been reported in venom isolated from *Pardosa pseudoannulata, Cyriopagopus hainanus* (formerly *Haplopelma hainanum), Selenocosmia jiafu, Lycosa singoriensis* and *Pamphobeteus verdolaga* (Zhang et al., 2010; Undheim et al., 2013; Cheng et al., 2016; Huang et al., 2018; Estrada-Gomez et al., 2019; Hu et al., 2019). The general assumption is that ICK peptides are highly diverse components of spider venom and dozens of different peptides may be present per species (Langenegger et al., 2019). In contrast, we found that ICK peptides were only a minor component of *A. bruennichi* venom, with only three different peptides identified (^~^5.5% of the overall diversity) and low abundance (only 3.3% of the total content based on TPM counts) suggesting a less important role in wasp spider venom compared to other spiders. Instead, we found that CAP superfamily proteins were both the most diverse (15 different members, >27% of the overall diversity) and the most abundant (>64% of the total content based on TPM counts), suggesting these proteins are particularly important for the function of *A. bruennichi* venom. Given that *A. ventricosus* venom also contains several CAP proteins (Duan et al., 2013), we speculate that CAP proteins may be generally important for venom function in orb weaving spiders (although we cannot compare our results directly to this previous study because it was based solely on proteomic analysis and lacked quantitative data). However, the atypical nature of *A. bruennichii* venom allows us to develop a functional hypothesis based on the ecology of wasp spiders, focusing on the most dominant venom components.

### 4.1 The importance of CAP superfamily proteins in wasp spider venom

The CAP superfamily is one of several protein groups that have undergone convergent recruitment and neofunctionalization in venom systems, and CAP proteins have therefore been isolated from the venoms of snakes, spiders, cone snails, scorpions, fish, cephalopods, and a variety of insects (Fry et al., 2009). This taxonomic ubiquity reflects the ability of CAP proteins to adopt diverse functions. For example, CAP proteins in snake venoms act as neurotoxins by interacting with ion channels (Brown et al., 1999; Yamazaki et al., 2002; Fry et al., 2009) whereas CAP proteins in the venoms of lampreys, hematophagous insects and ticks are thought to facilitate feeding (Ribeiro et al., 2004; Ito et al., 2007; Fry et al., 2009). In bees, wasps and ants, CAP proteins are major allergenic components of venom and are therefore associated with inflammation and potentially fatal anaphylaxis (Fang et al., 1988; Caruso et al., 2016). CAP proteins have been detected in several spider venoms but generally as minor components and their function is unknown (e.g. Undheim et al., 2013; Kuhn-Nentwig et al., 2019). Thus far, the only known spiders with venom dominated by CAP proteins are *A. ventricosus* (Duan et al., 2013) and *A. bruennichi*.

The CAP proteins in *A. bruennichi* venom, as in other arthropods, are unlikely to act as neurotoxins because they lack the C-terminal cysteine-rich domain that confers the neurotoxicity of CAP proteins in snake venom (Fry et al., 2009). Phylogenetic reconstructions indicate that spider CAP proteins are similar to Tex31, a well-characterized CAP protein from the venom of the cone snail *Conus textile* (Fry et al., 2009) that has proteolytic activity (Milne et al., 2003). This suggests that CAP proteins in *A. bruennichi* venom (and other spider venoms) may support extra-oral digestion or act as spreading factors to promote the uptake of other venom components. The lack of CAP superfamily neurotoxins in spiders would not be a disadvantage because the venom contains other neurotoxic components, including ICK peptides, prokineticins and MIT-atracotoxins. Assuming that the newly identified aranetoxins and neuropeptides also act as neurotoxins, wasp spider venom clearly contains an impressive arsenal of bioactive components that may facilitate hunting. Interestingly, an early study on effects of spider venom against cockroaches and meal beetles demonstrated that wasp spider venom can paralyze but not kill both these prey (Friedel and Nentwig, 1989). Therefore, despite the atypical composition of *A. bruennichi* venom, this species is nevertheless capable of neurotoxic envenomation.

### 4.2 Wasp spider venom contains potential new toxin classes similar to arthropod neuropeptides

Our proteomic analysis of *A. bruennichi* venom identified five polypeptides that we grouped as unclassified aranetoxins, showing high sequence similarity to the recently discovered U_6_ and U_8_ aranetoxins in *A. ventricosus* (Duan et al., 2013). They are not yet formally assigned to any known class of toxins and their molecular and biological functions remain to be determined. However, the five unclassified aranetoxins were expressed at high levels in our venom gland transcriptome dataset and are therefore likely to fulfill important functions in the *A. bruennichi* venom system. Their presence in two orb weavers but no other spider families suggests their role may be specific to the unique ecological niche of orb weavers.

We also identified one diuretic hormone-like peptide, one IGFBP and two ITG-like peptides. Diuretic hormone-like peptides contain a DH31-like domain and are related to diuretic hormone from the Florida carpenter ant (*Camponotus floridanus*) and, to a lesser extent, U-scoloptoxin-Sm2a from the centipede *Scolopendra morsitans* (Undheim et al., 2015). This class of protein has not been detected in other spider venoms. The *A. bruennichi* IGFBP is closely related to a protein found in the venom of the tiger wandering spider *Cupiennius salei* (Kuhn-Nentwig et al., 2011). Such proteins are commonly found in arachnid venoms but also in scorpions (*Superstitia donensis, Hadrurus spadix* and *Centruroides hentzi*) and ticks of the genus *Amblyomma* (Santibanez-Lopez et al., 2016; Esteves et al., 2017; Rokyta and Ward, 2017; Ward et al., 2017). A related protein is present in the venom of *A. ventricosus*, but its function has not yet been determined (Kono et al., 2019). Whereas the *A. bruennichi* diuretic hormone-like peptide and insulin-like growth factor-binding protein are relatively minor venom components, the two ITG-like peptides were expressed at high levels in the venom gland transcriptome and may therefore fulfil more important functions. They are closely related to peptides found in the black cutworm moth (*Agrotis ipsilon*) and *C. floridanus* but have not been identified in other spider venoms.

The role of all three classes of proteins described above is unclear, but hormones in other venomous animals have been weaponized as toxins to subdue prey. Such neofunctionalization might occur when hormones recruited to the venom gland for normal physiological activity undergo mutations that affect their surface chemistry and potential for functional interactions. For example, a neuropeptide that regulates physiological processes in the predator could become a toxin if a mutation causes it to interact with a different receptor in a prey species following envenomation. If this process occurs in the context of gene duplication and divergence, the new role in envenomation could be unlinked from the original physiological role allowing evolutionary forces to fix the neuropeptide as a venom toxin. The neofunctionalization of hormones and neuropeptides in venom systems is further highlighted by the recent discovery of the convergent recruitment of hyperglycemic hormones in the venom of spiders and centipedes (Undheim et al., 2015). This study demonstrated that helical arthropod-neuropeptide-derived (HAND) toxins are derived from hormones of the ion transport peptide/crustacean hyperglycemic hormone (ITP/CHH) family, which are ubiquitous and functionally diverse neuropeptides in arthropods. ITP/CHH peptides have also been recruited into the venom systems of ticks and wasps and are not restricted to the HAND family. For example, emerald jewel wasp (*Ampulex compressa*) venom contains tachykinin and corazonin neuropeptides that induce hypokinesia in cockroaches (Arvidson et al., 2019), whereas exendin from the venom of helodermatid lizards is a modified glucagon-like peptide that interferes with pancreatic insulin release (Yap and Misuan, 2019). Amphibians have also recruited a variety of hormone peptides as skin toxins (Gaudino et al., 1985; Roelants et al., 2010; Roelants et al., 2013; Lüddecke et al., 2018). The novel neuropeptides in *A. bruennichi* might fulfill other functions and their potential role as toxins must be tested, but the strong expression of the ITG-like peptides indicates an important function in the venom system.

### 4.3 The potential ecological role of atypical wasp spider venom

The atypical venom composition of *A. bruennichi* could be explained by trophic specialization, which would select for simple venoms prioritizing specific components needed to subdue selected prey species. This would contrast with generalist predators, where diverse venom components would confer a selective advantage (Daltry et al., 1996; Fry et al., 2003; Li et al., 2005; Phuong et al., 2016; Lyons et al., 2020). However, *A. bruennichi* is not regarded as a specialist feeder, and an alternative explanation must be sought (Szymkowiak et al., 2005).

In a pioneering study, the hunting behavior of three orb weavers (*Nephila claviceps, Argiope aurantia* and *Argiope argentata*) was compared using bombardier beetles (*Brachinus* spp.) as prey (Eisner and Dean, 1976). These beetles have evolved a unique chemical defense involving the stress-triggered release of phenolic compounds from abdominal glands under high pressure. The phenolic compounds undergo rapid exothermal oxidation to benzoquinones, thus spraying predators with a pressurized discharge at temperatures up to 100°C, allowing the beetle to escape from most situations (Dean et al., 1990). When such beetles were offered to the three spiders, the interactions were distinct: *N. claviceps* always tried to inject venom as a first attack strategy and this resulted in successful defense and escape by the beetles, whereas both *Argiope* species deployed silk as a first attack strategy and envenomation would follow only when the beetle was fully covered and unable to move (Eisner and Dean, 1976). In another study using *A. bruennichi* as the model predator, silk was deployed to overpower most prey insects, including those with other robust defense systems such as wasps, and only lepidopteran prey were attacked with venom first (Nyffeler and Benz, 1982). Similar findings have been reported for eight other *Argiope* species, suggesting that this specialized hunting behavior is highly conserved within the genus (Robinson, 1969; Harwood, 1974; Nyffeler and Benz, 1982). The prevalence of this silk-based hunting strategy may help to explain the simplicity of its venom, which would be under selection solely for its ability to subdue lepidopteran prey.

In venomous animals, each toxin is a valuable resource that contributes to fitness by facilitating predation, but this advantage must be balanced against the metabolic costs of replenishing venom stocks (Nisani et al., 2012; Saggiomo et al., 2017; Blennerhassett et al., 2019). Many venomous animals have evolved as trophic specialists to reduce these costs, and some even produce different venoms for different purposes (e.g. Walker et al., 2018). Spiders face a similar dilemma because they possess two potentially competing systems to subdue prey, namely their venom and silk glands. In both cases, protein resources are deployed as a means to facilitate predation, and in both cases the glands must be replenished at significant metabolic cost (Guehrs et al., 2008). We propose that the simplicity of *A. bruennichi* venom may reflect the evolutionary consequences of competition for resources between the venom and silk systems which have driven its behavioral specialization to use different strategies against different prey species. Intriguingly, the ‘silk-first’ strategy provides the wasp spider with the competitive advantage of a high success rate against even well defended prey (Eisner and Dean, 1976; Nyffeler and Benz, 1982) potentially contributing to its unprecedented success during its recent range expansion (Krehenwinkel and Tautz, 2013; Krehenwinkel et al., 2015).

## 5. Conclusions and future perspectives

Our detailed analysis of *A. bruennichi* venom identified potential new classes of toxins and potential new roles for known protein families including the predominant CAP superfamily. The molecular functions and biological roles of these proteins should be investigated in detail to disentangle the venom biology of wasp spiders and to identify new drug leads. The sequences we identified can be used to produce recombinant *A. bruennichi* venom proteins in larger quantities for detailed functional analysis, and that work is already underway in our laboratory. However, wasp spiders are small animals with limited venom yields, and are therefore unsuitable for traditional fractionation workflows (e.g. von Reumomont et al., 2014; von Reumont, 2018). Our venomics workflow overcomes this issue by combining the data-driven selection of interesting candidates based on venom gland transcriptome analysis with a dual proteomics strategy for the comprehensive identification of venom proteins directly. Importantly, our highly sensitive venomics workflow means that comprehensive venom composition is possible starting with only 14 spiders. A similar approach was recently used to analyze the venom proteome of the spider family Pholcidae (Zobel-Thropp et al., 2019).

Such novel workflows and technical platforms will help to extend our knowledge of venom composition beyond the small collection of amenable organisms with readily accessible venom systems. This has already shown that many commonly held assumptions about venom (based on the limited number of investigated species) are not supported when more diverse species are included. We describe the venom of *A. bruennichi* as atypical for spidersbecause its composition, dominated by CAP superfamily proteins and with ICK peptides fulfilling a minor role, differs from the restricted range of spider venoms that have been investigated thus far. Similarly, the recent proteomic analysis of pholcid venom also revealed a distinct composition dominated by neprilysin metalloproteases (Zobel-Thropp et al., 2019). Both studies highlight the importance of filling the taxonomic gaps in venom research in order to fully understand the hidden molecular diversity. This is likely to reveal there is no ‘typical’ spider venom but rather a spectrum of compositions reflecting different ecological niches. Such diversity will not only illuminate the field of arachnid evolutionary biology but will also provide many more promising candidates for translational research.

## 6. Acknowledgments

The authors are grateful to Philipp Heise, Lea Talmann and Anne Paas for their support in field and laboratory work. The authors acknowledge generous funding by the Hessen State Ministry of Higher Education, Research and the Arts (HMWK) for the project “Animal Venomics” via the LOEWE Center “Translational Biodiversity Genomics”.

## 6. Author contributions

TL, SL, AB, BMvR and AV designed the study. TL performed fieldwork and generated the sample material for transcriptome sequencing and venom proteomics. TL, SL and TT performed the laboratory proteomics work. TT and GL generated and analyzed the mass spectra. TL, FF, BMvR and AB generated and analyzed the transcriptome datasets. AV attracted the funding for the study. TL, BMvR SL wrote the manuscript with substantial input from AB, FF, GL, TT and AV.

